# Accurate and scalable demultiplexing of single-cell RNA sequencing using BEACON

**DOI:** 10.64898/2026.09.26.754667

**Authors:** Brian W. Ji, Marshall Lammers, Aubrey Houser, Nicholas Chan, Jose Rodriguez, Hannah Chae, Cheryl Xiang, Jack Bui, Hojun Li, Andrew L. Ji

## Abstract

Barcode-based multiplexing strategies can significantly increase sample throughput and decrease costs, while mitigating batch effects of single-cell RNA sequencing experiments. However, these approaches can be limited by inaccurate or inefficient demultiplexing, resulting in cell loss and reduced statistical power. Here, we present BEACON, a novel sample demultiplexing method that efficiently learns the background count distribution from data and removes it from individual cells, thereby improving classification accuracy. BEACON outperforms other state-of-the-art methods on multiple human data sets. We apply it to cancer cell line time course experiments in vitro, enabling the identification of genes associated with aggressive tumors in vivo. Finally, we adapt BEACON to multimodal protein-transcriptome profiling, enhancing protein signal recovery to identify a CD161-positive effector memory CD4 T-cell population with a Th17-like phenotype, which we prospectively validate. BEACON can therefore be applied to other droplet-based single-cell sequencing methodologies.

## Results

Sample barcoding is widely utilized for the multiplexing of single-cell RNA sequencing experiments. These approaches, which label cells with sample-specific barcodes, can increase sample throughput and decrease sequencing costs, while mitigating batch effects and removing multiplets^1–7^. Despite the development of numerous multiplexing strategies, sample demultiplexing, or the process of assigning cells to their original samples, remains a significant computational challenge^1,5,8–14^. This stems in part from the presence of background barcode reads inherent to droplet-based sequencing technologies that can fundamentally limit sample assignment accuracy^15^. Current methods can also scale poorly with large numbers of multiplexed samples. Indeed, when we attempted to demultiplex published single-cell transcriptomic datasets with existing methods, we found a high rate of classification failure or unrealistic multiplet assignments (**Supplementary Fig. 1**).

**Figure 1.**
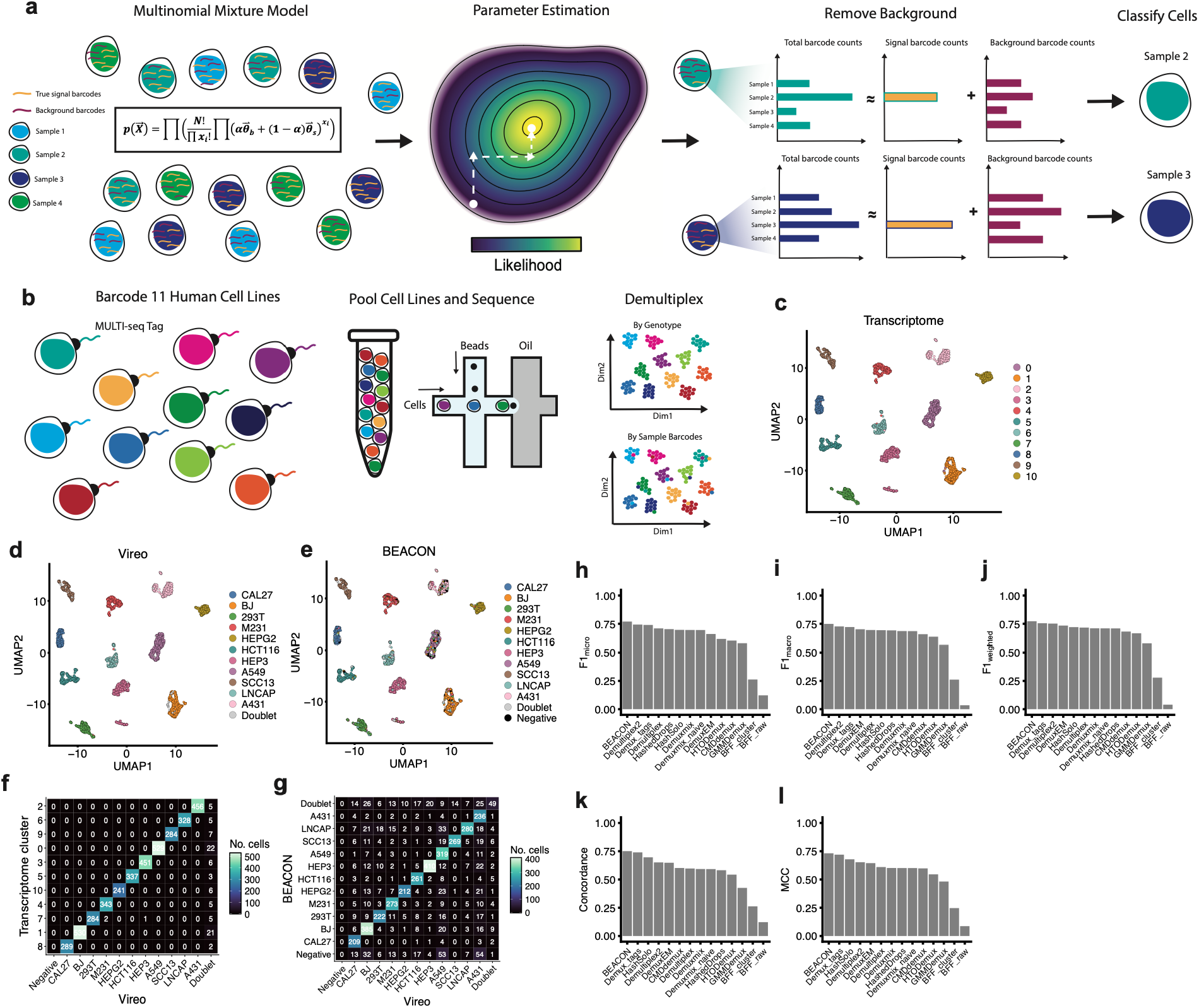
Accurate sample demultiplexing by BEACON on a panel of eleven human cell lines. **a)** Schematic of BEACON statistical model and classification workflow. **b)** Generation of an eleven human cell line ground-truth data set. Cell lines were barcoded using MULTI-seq, pooled, sequenced, and demultiplexed using SNP-based or barcode-based demultiplexing methods. **c)** UMAP representation of eleven human cell line transcriptomes. Cells are colored according to transcriptome clusters (Methods). **d)** As in **c)** but with cell line assignments colored using the SNP-based sample classification method Vireo. **e)** As in **c)** and **d)** but with cells colored based on sample assignments using BEACON. **f)** Joint distribution of Vireo and transcriptome cluster assignments across individual cells. **g)** Joint distribution of cell line classifications using Vireo and BEACON. **h-l)** Performance of barcode-based sample demultiplexing methods on the eleven cell line data set. Shown are **h)** F1_micro_, **i)** F1_macro_, **j)** F1_weighted_, **k)** concordance and **l)** Matthews Correlation Coefficient. Methods are ranked from highest to lowest in each panel. In **h-l)** Vireo assignments were used as ground truth.

To overcome these challenges, we have developed BEACON (Background Estimation and Assignment of Cell Oligonucleotides), a novel sample demultiplexing strategy that 1) learns the signal and background barcode count distributions directly from individual cells and 2) accurately classifies cells based on these learned distributions. While most methods rely on marginal barcode count distributions to make singlet, multiplet and negative sample assignments, BEACON fits a multinomial mixture model to the joint distribution of barcode counts across the entire dataset. Parameter estimation on this high-dimensional landscape would typically not be practical. We therefore derived an efficient, iterative algorithm to climb the joint-likelihood function (**Fig. 1a**, Methods). We carried out a number of analyses to ensure the robustness of our approach. First, we confirmed using Monte Carlo simulations the ability of BEACON to accurately recover known parameters across input data of widely differing complexity, and across a range of method input parameters (**Supplementary Fig. 2**). Second, we confirmed fast parameter convergence of our iterative likelihood-climbing algorithm (**Supplementary Fig. 3a-d**). Third, we confirmed that data reconstructed by BEACON is highly correlated with original count data across individual datasets, supporting the validity of the multinomial mixture model (**Supplementary Fig. 3e**,**f**). Lastly, for data sets split into technical sequencing replicates that share the same underlying background read count distribution, we confirmed that the estimated background distributions learned by BEACON were highly similar (**Supplementary Fig. 3g-i**).

**Figure 2.**
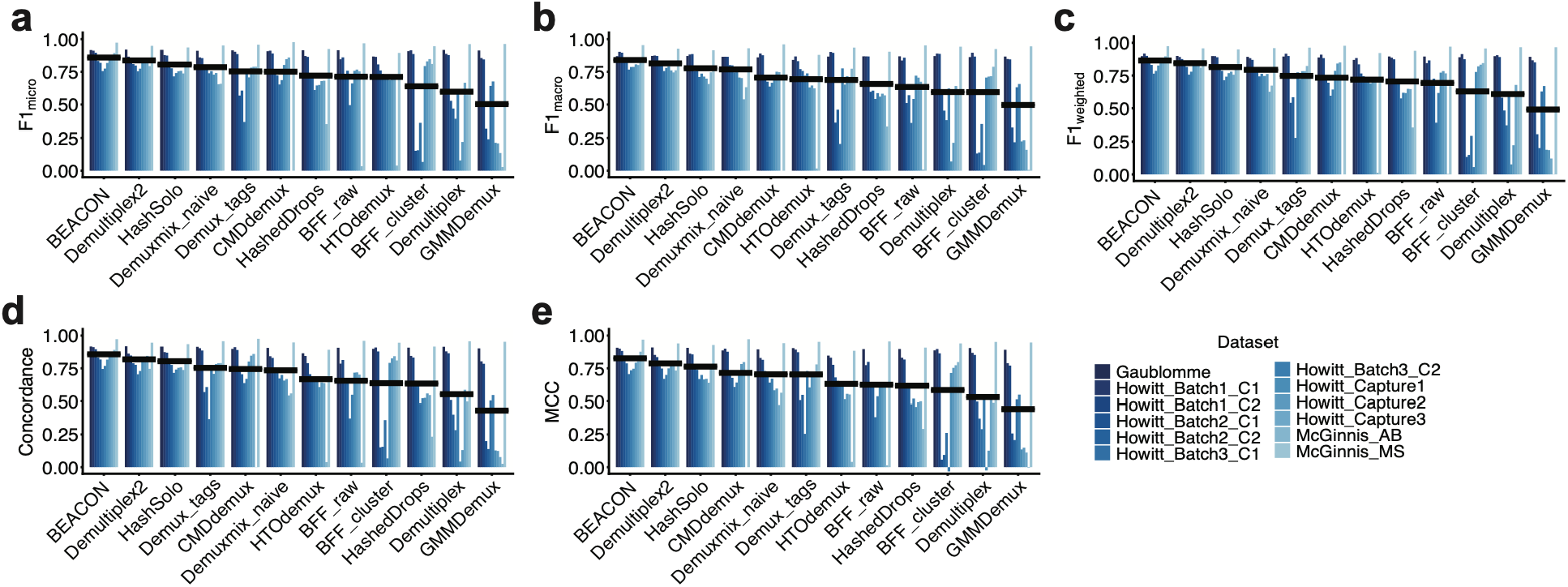
Performance of BEACON across multiple human data sets. Shown are **a)** F1_micro_, **b)** F1_macro_, **c)** F1_weighted_, **d)** concordance and **e)** Matthews Correlation Coefficient of various barcode-based demultiplexing strategies applied to twelve human data sets. Individually barcoded samples within each dataset were also genetically distinct. Within each panel, methods are ranked by average scores across the data sets, indicated by black horizontal lines. If a method was unable to run, it was assigned a score of zero for that particular data set. Genotype-based assignments were used as ground truth.

**Figure 3.**
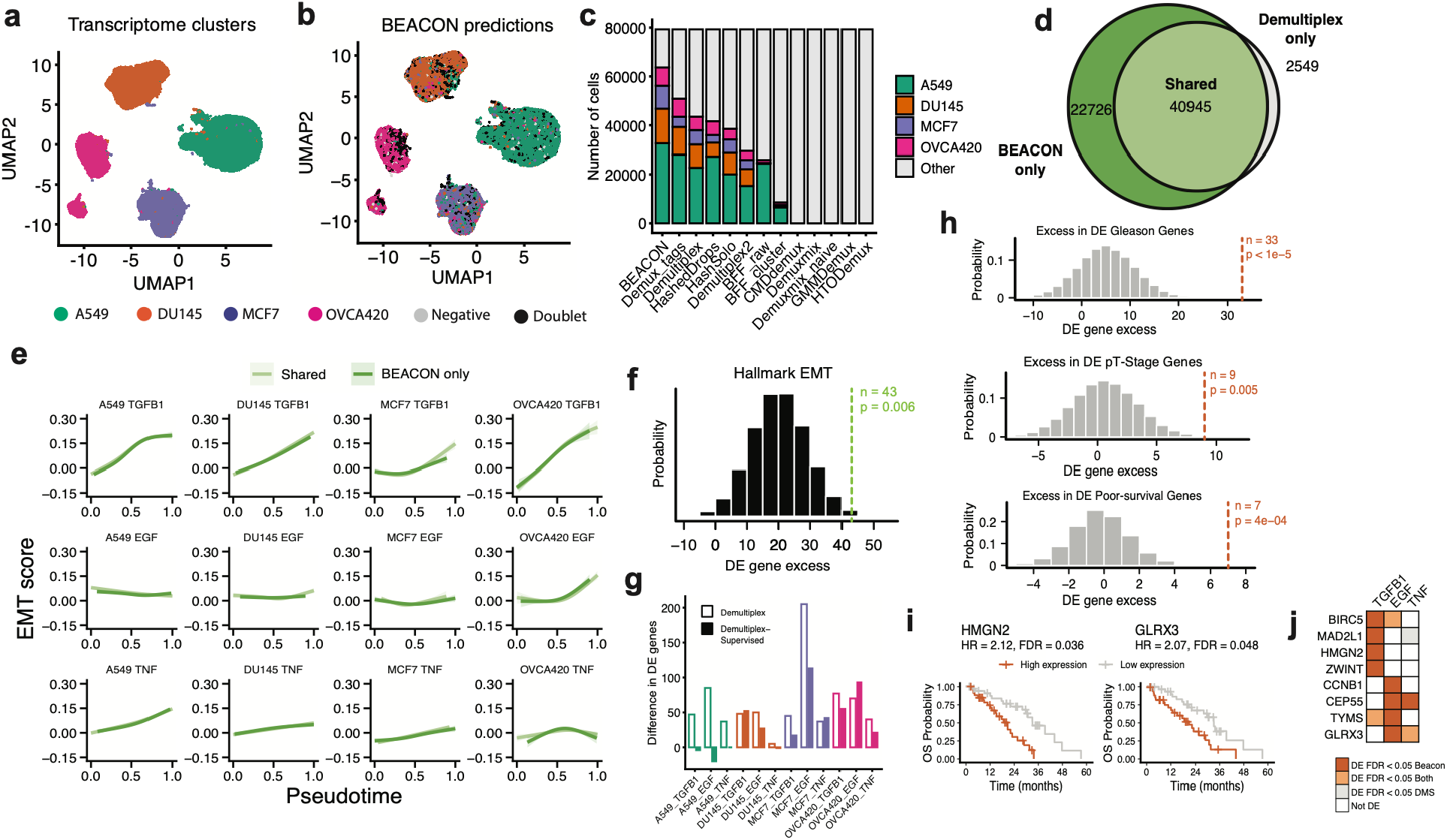
Application of BEACON to human cancer cells treated with EMT-promoting ligands. **a)** UMAP representation of 79,312 single-cell transcriptomes from the study of Cook and Vanderhyden. Cells are colored based on transcriptome cluster and annotated by inferred cell line identity. **b)** As in **a)** but with colors representing cell line identities predicted by BEACON. **c)** Concordance between transcriptome-based and sample demultiplexing-based cell line identities, stratified by method and cell line. Methods are ranked by total number of concordant cell line predictions. Gray bars denote cells with discordant transcriptome-based and sample demultiplexing-based predictions. These by default include doublet and negative demultiplexing annotations. Stacked bars sum to the total number of sequenced cells in the dataset. **d)** Venn diagram of concordant transcriptome-based and barcode-based cell line identities, using BEACON and Demultiplex. **e)** EMT gene scores as a function of pseudotime for each of the twelve time series experiments, using cell subsets as indicated in **d)**. Smoothed GLM fits are shown for each time series. No significant differences in gene score dynamics were observed between any of the twelve pairs by permutation testing (Methods). **f)** Difference in differentially expressed EMT genes identified by BEACON as compared to Demultiplex across the entire dataset, indicated by the green vertical line. Histogram represents the null distribution generated from 1 million random subsamplings of equal gene size. One-sided permutation-based p-value is indicated (Methods). **g)** Difference in the number of differentially expressed genes identified by BEACON compared to both Demultiplex and Demultiplex-supervised for each of the twelve time series. **h)** Excess in differentially expressed (DE) genes identified by BEACON compared to Demultiplex-supervised that overlap with genes positively associated with Gleason score and pathologic T stage in primary prostate adenocarcinoma tumors, and poor overall survival in metastatic castrateresistant prostate tumors. **i)** Kaplan-Meier survival curves of high and low-expressing tumors for HMGN2 and GLRX3. **j)** Genes identified by BEACON as differentially induced by the indicated ligands that are associated with poor overall survival in advanced prostate cancer.

We next sought to benchmark the performance of BEACON using data where the underlying sample assignments have been established. Demultiplexing strategies based on natural genetic variation can provide an accurate orthogonal means to classify samples, provided that each sample is genetically distinct^16–20^. To create such a “ground-truth” data set, we pooled eleven different human cell lines uniquely tagged using MULTI-seq barcodes^1^, and sequenced the resulting cell line mixture (**Fig. 1b**). As expected, we identified eleven distinct transcriptome clusters (**Fig. 1c**). Importantly, when classifying cells using the genotype-based demultiplexing method Vireo^16^, we found near-perfect concordance between cell line annotations and transcriptome clusters, supporting genotype-based demultiplexing as a reliable ground truth reference (**Fig. 1d,f,** Methods).

We next used BEACON to make sample classifications, finding overall agreement with Vireo assignments (**Fig. 1e,g)**. To facilitate quantitative comparisons between multiple demultiplexing methodologies, we calculated five metrics of classification performance that balance inherent tradeoffs in the data^21^: F1_micro_ score, F1_macro_ score, F1_weighted_ score, Matthews Correlation Coefficient (MCC), and overall concordance (Methods). We found BEACON outperformed all other methods using each of the five metrics (**Fig. 1h-l**).

To further assess the performance of BEACON, we compiled a list of twelve additional human data sets, spanning multiple publications and sample barcoding strategies^5,15,22^. These datasets were specifically selected because barcoded samples were also genetically distinct, facilitating concomitant “ground truth” assignments via SNP-based classification. We again found that BEACON outperformed all other methods using each of the five metrics, when averaged across all data sets (**Fig. 2**). We also found, with few exceptions, that BEACON tended to perform superiorly on individual datasets (**Supplementary Fig. 4**) and at the individual sample level (**Supplementary Fig. 5**). We observed similar results when downsampling total barcode counts from each dataset by an order of magnitude (**Supplementary Fig. 6a**). Importantly, we found that the performance of BEACON was robust across a range of input parameters (**Supplementary Fig. 6b**,**c**).

**Figure 4.**
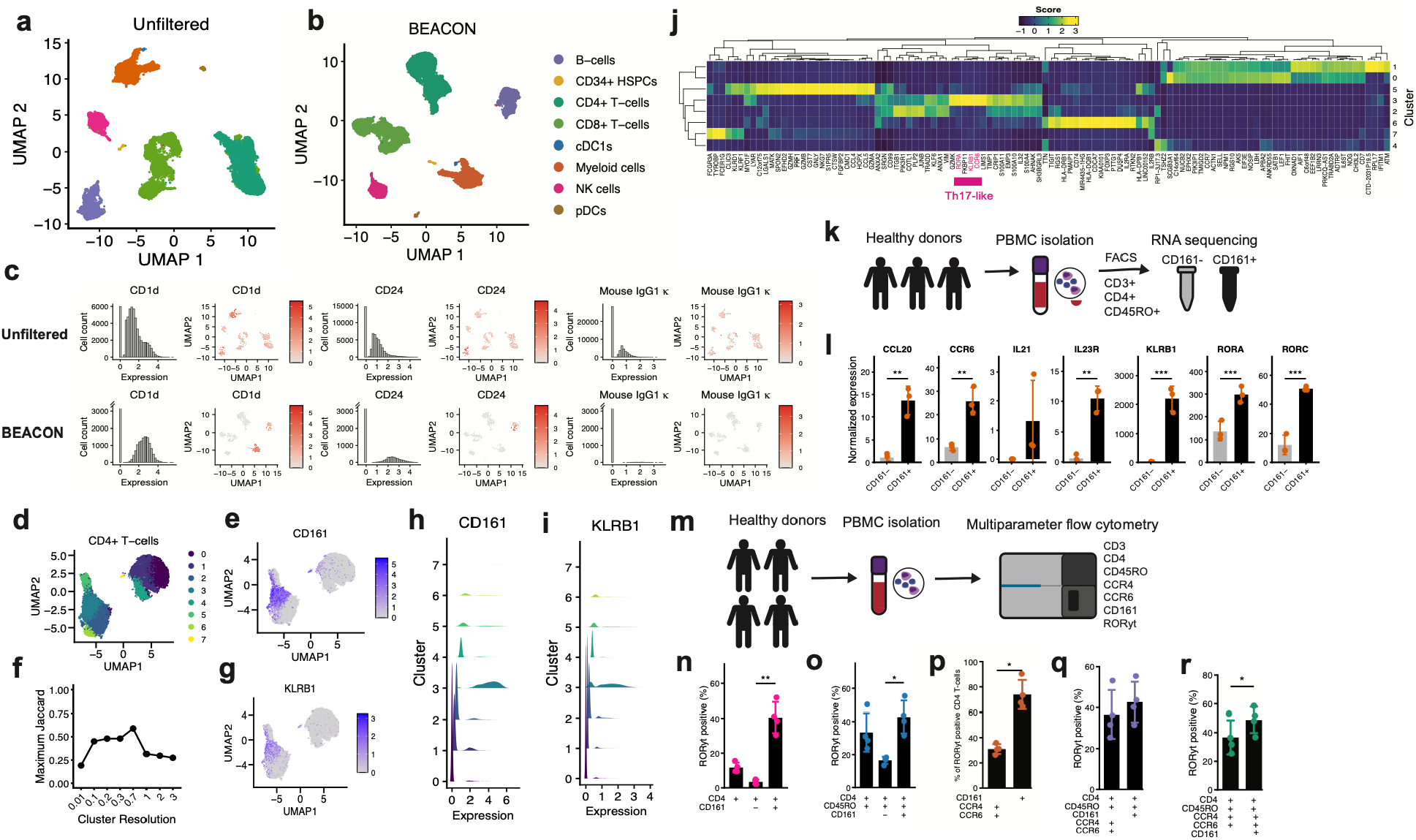
BEACON model adapted to CITE-seq of human PBMCs. **a**,**b)** UMAP representation of 53,846 cells from the study of Kotliarov et al., based on CITE-seq measurements of 82 cell surface protein markers. Unfiltered ADT abundances are shown in **a)**, while BEACON-filtered ADT abundances are shown in **b). c)** Expression of select protein abundances as measured by CITE-seq using unfiltered and BEACON-filtered data. **d)** CD4+ T-cells clustered using BEACON-filtered ADT abundances. **e)** BEACON-filtered CD161 ADT expression of CD4+ T-cells. **f)** Maximum Jaccard index between the CD161-high cluster in **d)** and clusters in the unfiltered CITE-seq data using various Louvain clustering resolution parameters. **g)** KLRB1 RNA expression of CD4+ T-cells overlaid on the UMAP from **d). h**,**i)** Histograms of BEACON-filtered CD161 expression and KLRB1 RNA expression within clusters. Histograms are scaled by the total number of cells in each cluster. **j)** Heatmap of genes whose expression is significantly enriched within individual clusters (Methods). Genes associated with the Th17 lineage are highlighted in pink. **k)** Prospective isolation of CD161-positive and CD161-negative populations from the peripheral blood of three healthy human donors subjected to bulk RNA-seq. **l)** Expression of canonical Th17- associated genes in the CD161-positive and CD161-negative populations. Only genes detected in more than a single sample were included. Asterisks indicate significance as assessed by the DESeq2 Wald test using a paired design that accounts for donor identity and tests the effect of CD161 status. **m)** Multiparameter flow cytometry of the indicated surface and intracellular proteins from the PBMCs of four healthy human donors. **n-o)** RORγt-positivity within the indicated CD4 compartments. **p)** Percentage of CD161-positive or CCR4/CCR6 double-positive cells among RORγt-positive CD4 T-cells. **q-r)** RORγt-positivity within the indicated CD4 T-cell subsets. *p < 0.05, **p<0.01, *** p<0.001.

We then asked how well BEACON would perform on datasets with higher numbers of samples. We therefore analyzed the data from Cook and Vanderhyden, who multiplexed 96 sample conditions comprising twelve time course experiments of four cancer cell lines (A549, DU145, MCF7, OVCA420) each treated with three ligands (TGFB, EGF, TNFa) previously shown to promote the epithelial-to-mesenchymal transition (EMT)^23^. Consistent with the original publication, we found that major transcriptome clusters mapped to individual cell lines (**Fig. 3a**). As ground truth assignments were not available in this data, we reasoned that cell line concordance rates could serve as a proxy for classification performance, assuming doublet rates in the data are relatively low, and that sample misclassification is equivalent within and between different cell lines. We found that BEACON predictions largely agreed with transcriptome-based assignments (**Fig. 3b**). Strikingly, BEACON assigned roughly 10,000 more cells than the next-best-performing method, while maintaining a low rate of discordant cell line predictions (**Fig. 3c, Supplementary Fig. 7**). We were surprised to find that many methods failed to make any predictions at all on this data set.

Several additional lines of evidence support the accurate recovery of cells by BEACON. First, when compared to Demultiplex, the method used in the original publication, EMT expression dynamics in cells assigned to their corresponding transcriptomic clusters by BEACON alone were highly similar to those in cells concordantly assigned by both methods (**Fig. 3d,e**), as well as to patterns reported in the original publication (**Supplementary Fig. 8a-f**). Second, differentially expressed genes using cells identified by BEACON alone were enriched for EMT involvement (**Supplementary Fig. 8g**). Third, cells assigned to their corresponding transcriptome cluster by BEACON frequently shared a consistent assignment with at least one additional method, although BEACON also uniquely assigned the most cells to their corresponding transcriptome cluster (**Supplementary Fig. 8h-j**). Collectively, these results support accurate demultiplexing and increased cell recovery rates using BEACON, even in data sets with a large number of multiplexed samples.

We hypothesized that BEACON could increase the power to detect differentially expressed genes in this data. Indeed, when compared to Demultiplex, BEACON identified a greater overall number of differentially expressed genes with significant enrichment for EMT-related genes, supporting the recovery of EMT gene expression dynamics previously underpowered for detection (p=0.006, Methods, **Fig. 3f**). As we attempted to compare the identities of differentially expressed genes with the original publication, we discovered the authors used a modified implementation of Demultiplex (hereafter referred to as Demultiplex-supervised), whereby select samples from individual sequencing pools were manually omitted to counterintuitively improve singlet recovery. Comparing BEACON to both Demultiplex and Demultiplex-supervised, we again found an overall increase in the number of significant differentially expressed genes (**Fig. 3g, Supplementary Fig. 8k**).

In light of these findings, we reasoned that the genes identified as differentially expressed by BEACON might also be reflected in aggressive tumor phenotypes in vivo. Using prostate cancer as an example, we first identified genes independently associated with either pathologic T stage or Gleason scores from primary prostate adenocarcinoma tumors^24^ (Methods). Within these sets of genes, we then quantified the excess number of differentially expressed genes identified by BEACON relative to Demultiplex-supervised in DU145 prostate cancer cells (Methods). Notably, in both cases, this excess was significantly greater than expected by chance (**Fig. 3h, Supplementary Fig. 9a**,**b**), suggesting that BEACON improves the ability to identify genes that are specifically associated with clinically aggressive phenotypes in primary patient tumors. To further test this idea, we analyzed an independent cohort of metastatic castrate-resistant prostate cancer (mCRPC) to identify genes that are associated with poor overall survival^25^ (Methods). Indeed, among this set of genes, BEACON identified a significantly higher number of differentially expressed genes (**Fig. 3h**). In total, BEACON uniquely nominated eight genes whose expression was significantly associated with poorer overall survival in patients with mCRPC (**Fig. 3i**,**j, Supplementary Fig. 9c**). Some of these genes, such as CEP55, CCNB1, and BIRC5, have been previously linked to aggressive forms of prostate cancer^26–28^. Others, such as GLRX3 and HMGN2, have not been previously characterized in prostate cancer and represent potential candidates for further investigation.

Finally, we sought to generalize the BEACON framework to enhance signal recovery in other droplet-based sequencing modalities. To that end, we adapted the BEACON model to CITE-seq and applied it to a dataset of peripheral blood mononuclear cells (PBMCs) from 20 healthy human donors, in which single-cell transcriptomes were jointly profiled with 82 cell surface proteins using antibody-derived tags (ADTs) (Methods)^2,29^. Consistent with previous reports, background removal did not substantially affect major immune cell populations identified using ADT expression alone^30^ (**Fig. 4a**,**b, Supplementary Fig. 10a**). However, BEACON did reveal the underlying cell-type-specific expression patterns of several cell surface protein markers such as CD1d and CD24, while effectively eliminating signal from isotype controls (**Fig. 4c, Supplementary Fig.10b**). We next wondered if BEACON could identify more non-discrete immune cell subpopulations. Interestingly, when focusing on CD4+ T-cells, we identified an antigen-experienced cell population characterized by high CD161 expression that was not readily identifiable using either the unfiltered ADT or transcriptome data, regardless of clustering resolution (**Fig. 4d-f**,**h, Supplementary Fig. 10c-g**). This cluster was shared across all donors in the dataset and demonstrated significantly increased KLRB1 expression, the gene encoding CD161 (**Fig. 4g**,**i, Supplementary Fig. 10h-k**). CD161 is a C-type lectin receptor best known as a marker of the Th17 lineage^31,32^. Indeed, transcriptome analysis further identified RORA and CCR6, canonical markers of the Th17 lineage, and RORC, the master Th17 transcriptional regulator, as significantly and specifically expressed in this subpopulation (**Fig. 4j, Supplementary Fig. 10i**).

To determine whether this population could be prospectively isolated, we collected PBMCs from three healthy human donors and sorted CD161-positive and CD161-negative cells within the CD4-positive, CD45RO-positive compartment (**Fig. 4k, Supplementary Fig. 11**). Differential gene-expression analysis demonstrated increased expression of canonical Th17-associated genes in the CD161-positive population, further supporting CD161 as a marker that enriches for this phenotype, as well as genes associated with effector T-cell functions (**Fig. 4l, Supplementary Fig. 12**). We next asked whether CD161 also enriches for RORγt, the master Th17 transcriptional regulator encoded by RORC, at the protein level. We therefore collected PBMCs from four healthy human donors and performed multiparameter flow cytometry with surface staining for CD161 and intracellular staining for RORγt (**Fig. 4m**). RORγt-positive cells were significantly enriched within CD161-positive cells relative to CD161-negative cells, confirming findings from the CITE-seq and bulk RNA-seq experiments (**Fig. 4n**,**o**). Finally, we compared CD161 with CCR4 and CCR6, two additional surface markers commonly associated with the Th17 phenotype^33^. Among RORγt-positive CD4 T-cells, CD161 identified a significantly larger fraction of cells than the CCR4/CCR6 double-positive phenotype (**Fig. 4p**). Conversely, CD161-positive cells showed a trend toward greater enrichment for RORγt-positivity than CCR4/CCR6 double-positive cells. Further restricting the CCR4/CCR6 double-positive compartment to CD161-positive cells increased the proportion of RORγt-positive cells (**Fig. 4q- r**). Together, these findings establish CD161 as an effective surface marker that enriches for Th17-associated CD4 T-cells alone and in combination with CCR4 and CCR6. More broadly, these results demonstrate that BEACON can improve recovery of informative protein measurements in single-cell multimodal RNA-protein datasets, thereby revealing biologically meaningful cell populations that are obscured by ambient background counts.

## Discussion

Here, we present BEACON, a novel, scalable and robust sample demultiplexing strategy for single-cell RNA-sequencing studies. We demonstrate the superior performance of BEACON on multiple data sets and generate an original, validated ground-truth data set that will provide a useful resource for future method development and benchmarking. By improving cell recovery and assignment accuracy, BEACON increases the power to detect differentially expressed genes and to resolve gene expression dynamics. When applied to prostate cancer cell lines treated with EMT-promoting ligands in vitro, BEACON specifically revealed genes associated with aggressive tumor phenotypes and poor clinical outcomes in vivo, highlighting the importance of accurate cell assignment to uncover clinically relevant transcriptional programs. Along these lines, we anticipate that BEACON may also improve the ability to detect rare cell states or tumor populations in vivo and their underlying gene programs from sparse single-cell sequencing data. Finally, we generalize BEACON to enhance signal recovery in multimodal protein-RNA single-cell data sets to identify a CD4-positive, CD161-positive Th17-like effector memory T-cell population, which we prospectively isolate and validate using multiparameter flow cytometry and FACS. These results highlight the potential for BEACON to identify biologically relevant cell populations across various disease contexts that would otherwise be obscured by background read counts using conventional analysis. We further anticipate that the BEACON framework may also be broadly applicable to other multimodal single-cell profiling technologies^34,35^, wherein background count contamination represents an inherent challenge. Collectively, our results highlight the importance of novel analytic frameworks to harness the full potential of rapidly expanding singlecell genomic technologies.

## Author contributions

Conceptualization: B.J., H.L., A.J. Method development: B.J. Multiplexing and single-cell library preparation: A.H. Flow cytometry and FACS: M.L. Data analysis: B.J., M.L., A.H., N.C., J.R., C.X., J.B., H.L., A.J. Writing – original draft: B.J. Writing – review and editing: B.J., M.L., H.L., A.J. Supervision: H.L., A.J.

