## supplementary_figures for "Accurate and scalable demultiplexing of single-cell RNA sequencing using BEACON"

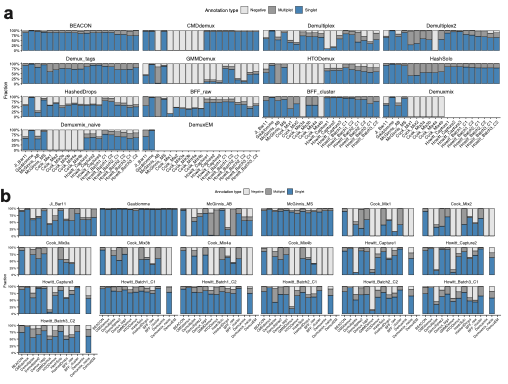

**Supplementary Figure 1. Annotation types across multiple datasets and demultiplexing methods**. **a)** Droplet annotation types are categorized as singlets, multiplets, or negatives, stratified by sample demultiplexing method. Methods that resulted in an error or did not finish on a particular dataset were by default assigned all negatives. If data were not available to run a particular method, the corresponding barplot was omitted. **b)** Annotation types as in **a)** but stratified by dataset.

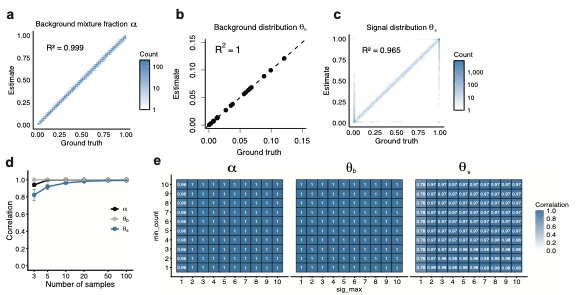

**Supplementary Figure 2. BEACON parameter estimation on simulated data. a-c)** Correlation between ground truth and BEACON-estimated parameters for an example simulated barcode count matrix ($m=20$ barcodes, $n=10,000$ cells) (Methods). Shown are the ground truth and recovered parameters using the BEACON algorithm for **a)** background count fraction $\alpha$, **b)** background barcode distribution $\theta_{b}$, and **c)** signal barcode distribution $\theta_{s}$. **d)** BEACON parameter recovery as a function of sample multiplexing complexity $m$. For a given sample complexity, ten random barcode count matrices were generated, where for each iteration we randomized the number of cells $n$, the background fraction distribution $\alpha$, the background barcode distribution $\theta_{b}$, and the signal barcode distribution $\theta_{s}$. Shown are the mean and standard deviation of correlations between the ground truth and recovered parameters. **e)** BEACON parameter recovery as a function of input parameters. We scanned a range of input sig_max values and min_count values as inputs to BEACON. Shown are correlations between recovered and ground truth parameters at a sample complexity of $m=30$ samples.

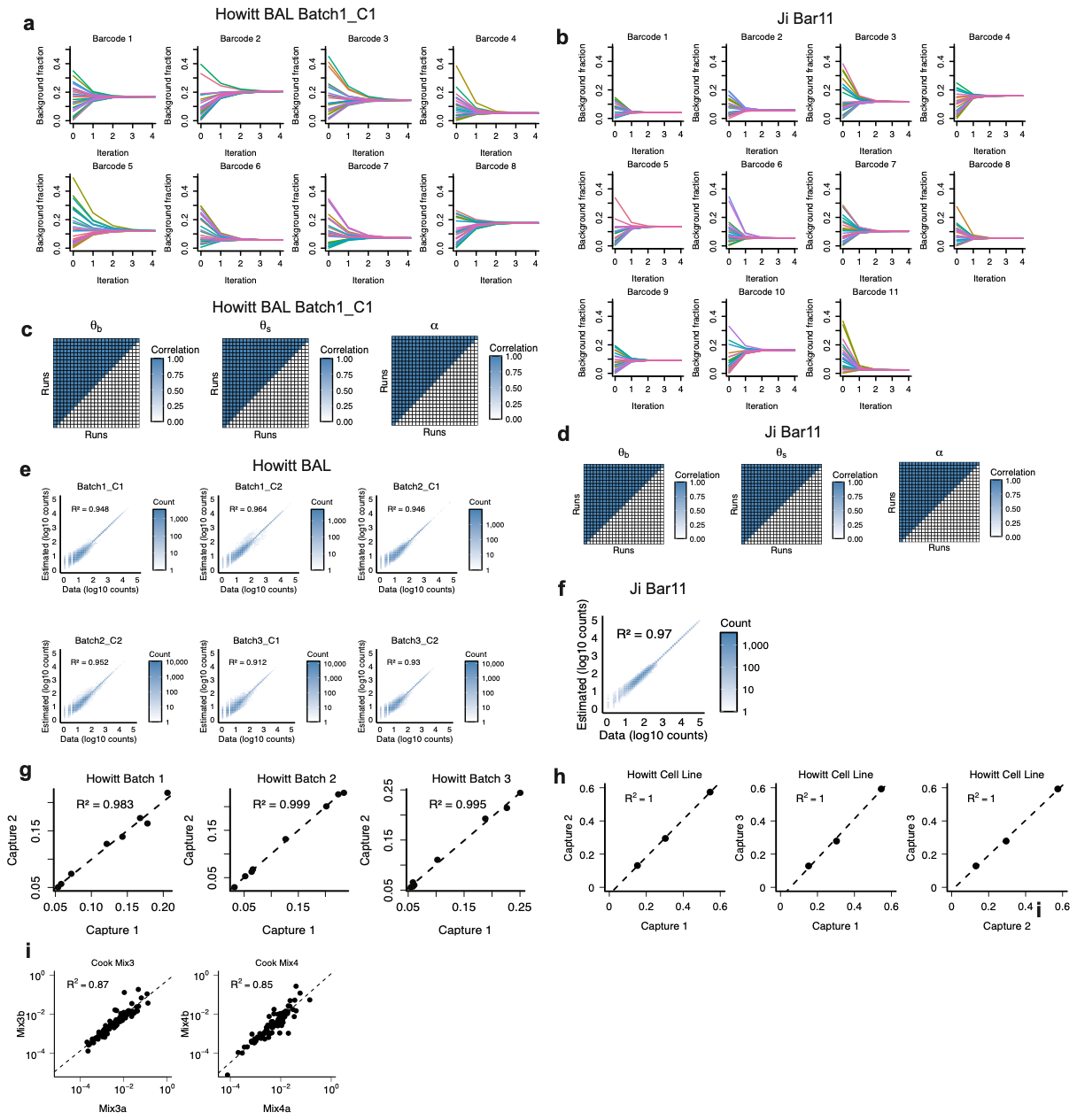

**Supplementary Figure 3. BEACON convergence on multiplexed datasets. a,b)** Convergence of each of the estimated background distribution fractions as a function of BEACON iteration across 20 random parameter initializations on the Howitt BAL and Ji Bar 11 datasets. **c,d)** Correlation of final parameters across 20 random parameter initializations from the Howitt BAL and Ji Bar 11 datasets. **e-f)** Correlation between estimated barcode counts reconstructed from BEACON and observed barcode counts for the Howitt BAL and Ji Bar 11 datasets**.** Correlation between estimated background distributions between technical replicates in the **g)** Howitt BAL dataset, **h)** Howitt cell line dataset, and **i)** Cook dataset.

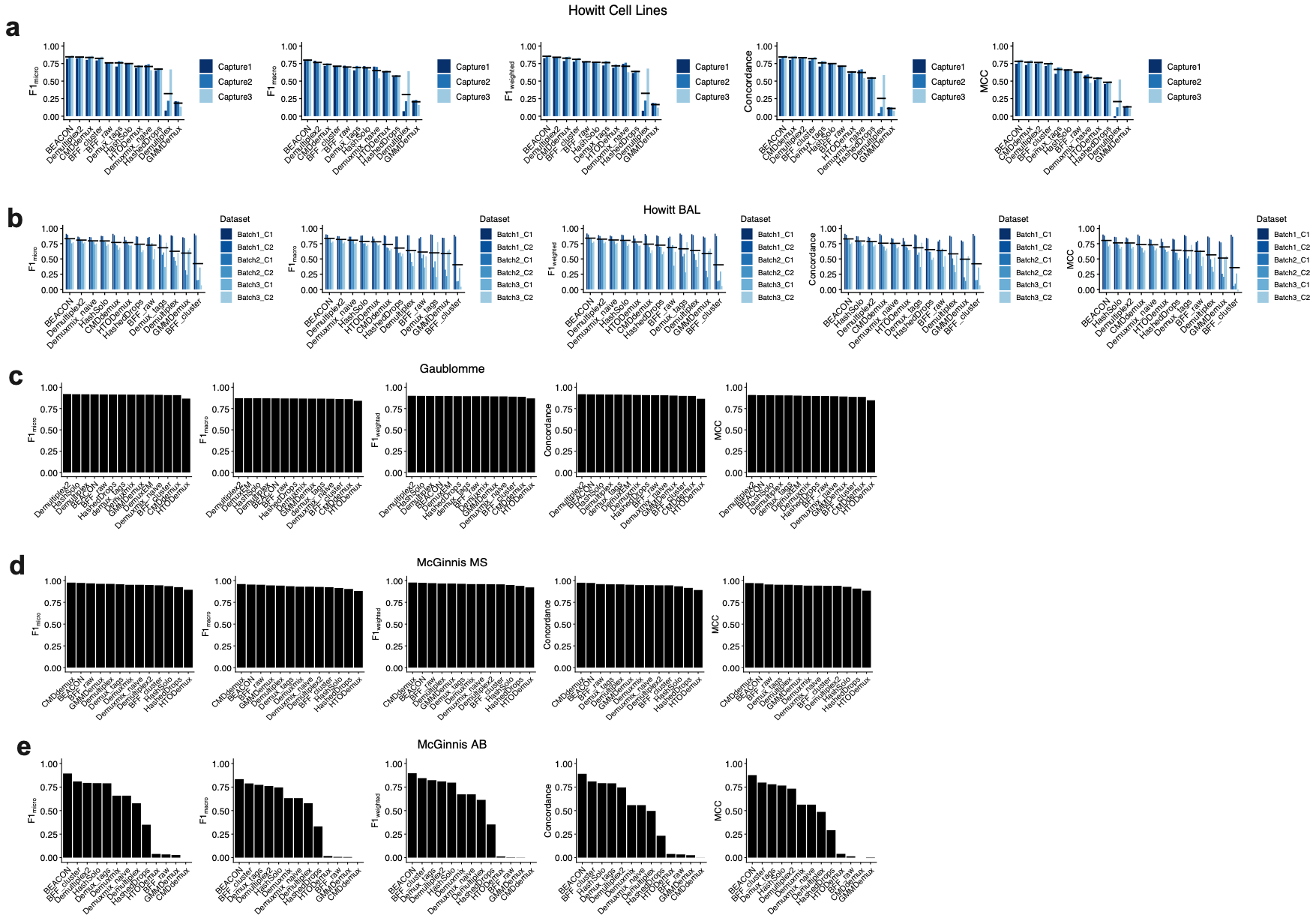

**Supplementary Figure 4. BEACON performance across multiple human data sets.** F1_micro_, F1_macro_, F1_weighted_, concordance and Matthews Correlation Coefficient (MCC) were calculated for different sample demultiplexing methods applied to multiple ground truth data sets as in Figure 2 of the main text. Within each panel, now stratified by individual data set, methods are ranked in descending order. Shown are data from the datasets of **a)** Howitt cell line, **b)** Howitt BAL, **c)** Gaublomme, **d)** McGinnis_MS, **e)** McGinnis_AB.

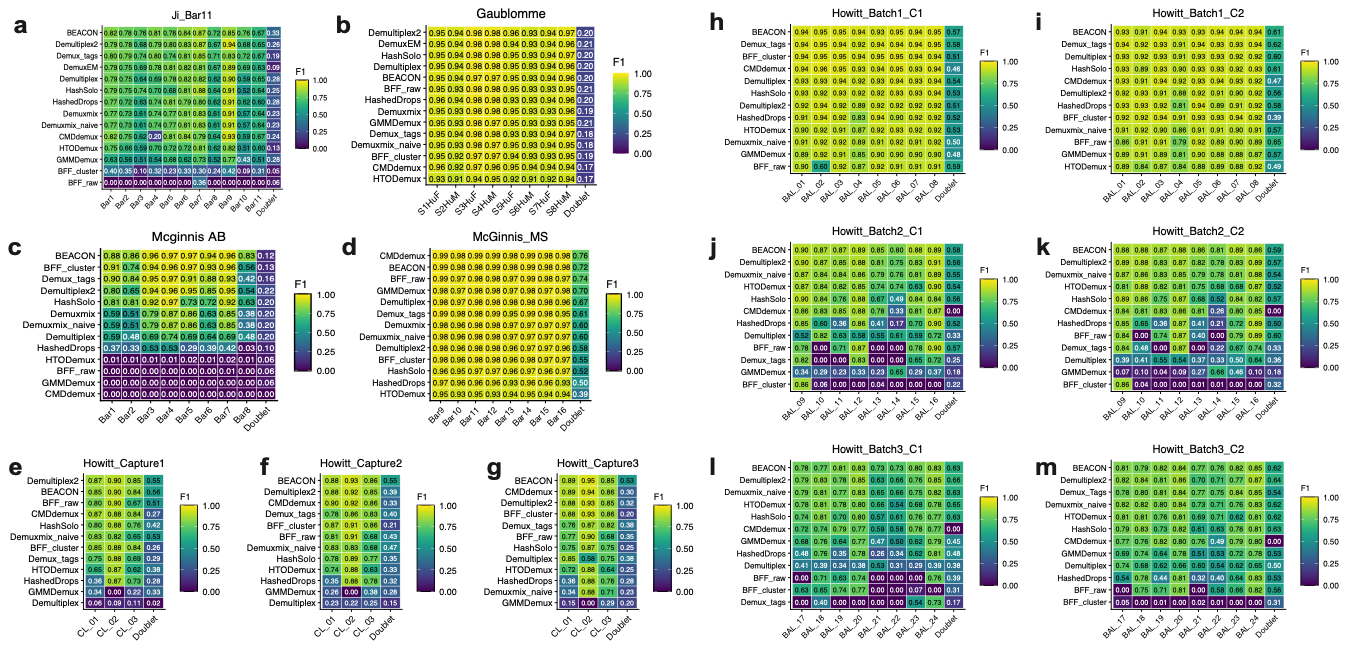

**Supplementary Figure 5. Classification performance at the individual sample level.** Individual F1 scores were calculated for each sample barcode, using the indicated methods and data sets. Shown are data from **a)** Ji_Bar11, **b)** Gaublomme, **c)** McGinnis_AB, **d)** McGinnis_MS, **e-g)** Howitt cell line and **h-m)** Howitt BAL. For each dataset, methods are ranked from top to bottom by average F1 score across barcodes.

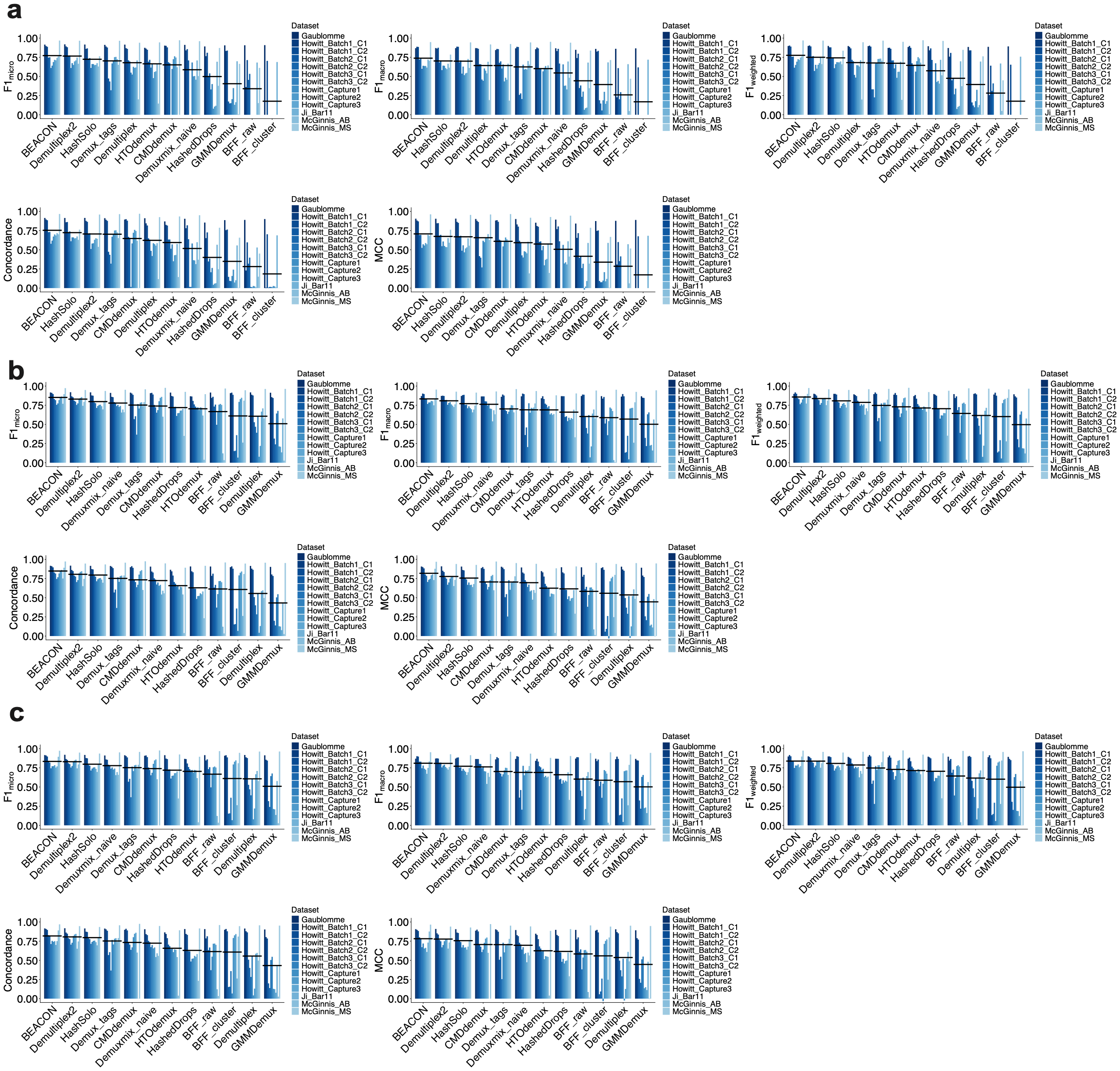

**Supplementary Figure 6. BEACON performance on downsampled data and across variation in input parameters.**  **a)** For each of the indicated datasets, barcode counts were downsampled 10-fold within individual cells. F1_micro_, F1_macro_, F1_weighted_, concordance and Matthews Correlation Coefficient (MCC) were then calculated. For each metric, methods are ranked by average value across all data sets. **b,c)** Classification performance when deviating BEACON input parameters from default values (Methods): **b)** sig_max = 2 (default 3), **c)** min_count = 15 (default 2).

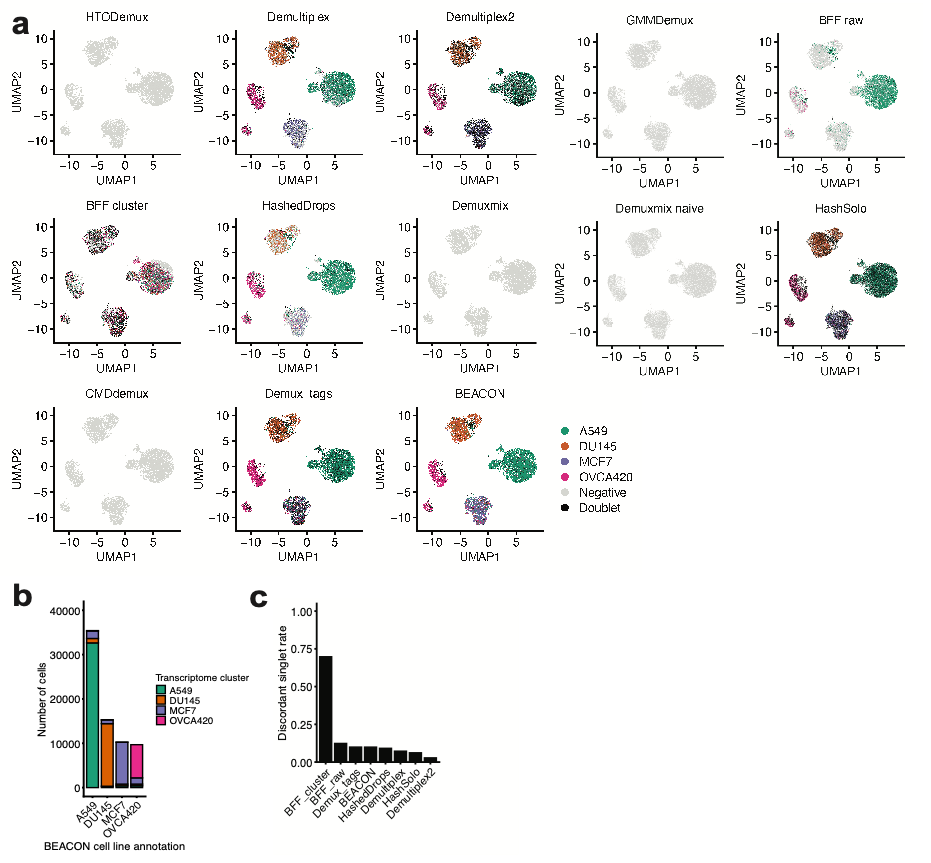

**Supplementary Figure 7. Cell line predictions on the Cook data set. a)** Cell line predictions overlaid on single-cell transcriptomes from the data of Cook and Vanderhyden across various sample demultiplexing strategies. **b)** Distribution of transcriptome clusters within each BEACON cell line annotation. **c)** Rates of discordant cell line predictions as a proportion of all singlet predictions for each of the sample demultiplexing strategies. CMDdemux, Demuxmix, Demuxmix_naive and GMMDemux did not predict any singlets and therefore rates of discordant cell line predictions could not be reported.

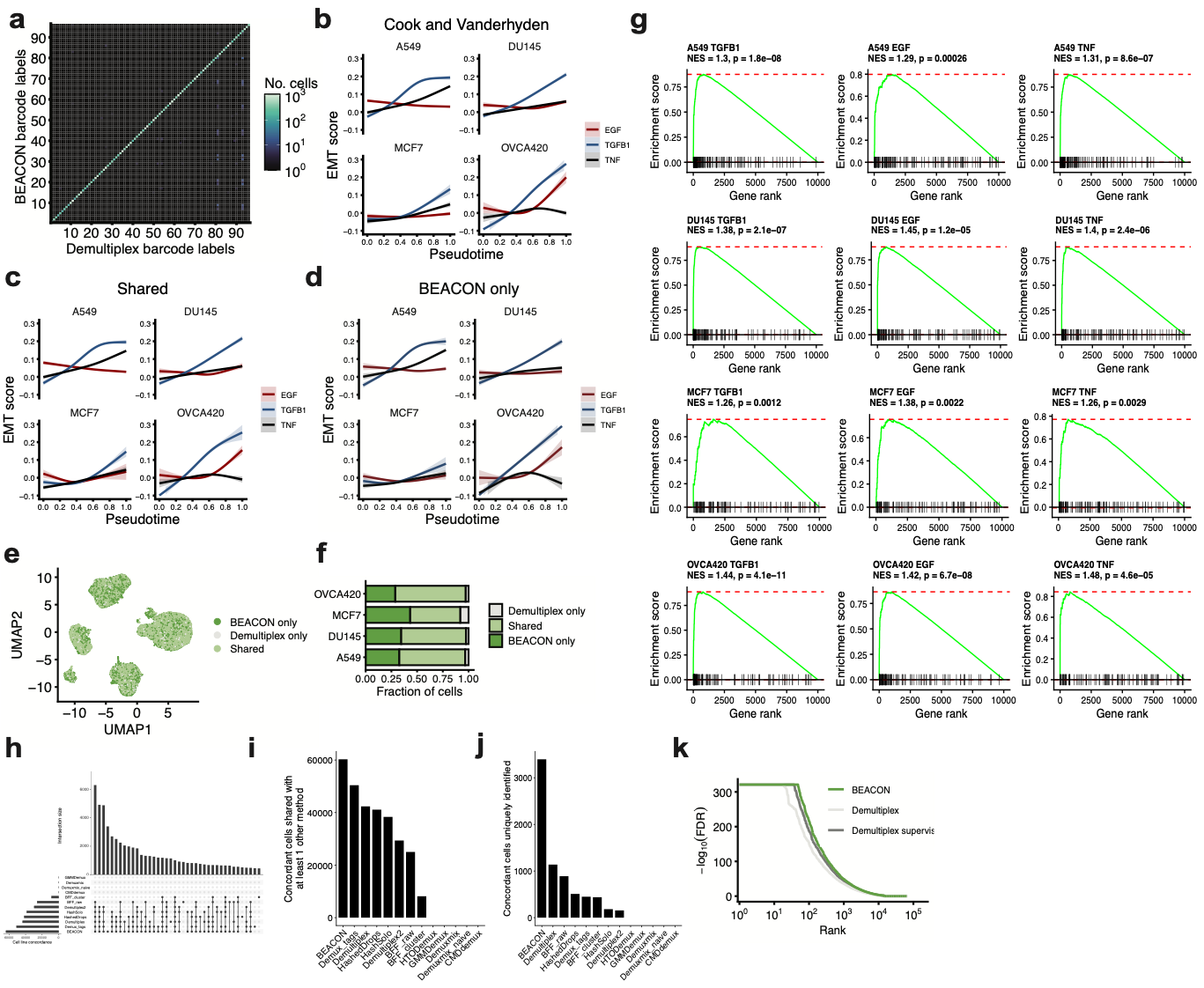

**Supplementary Figure 8. Improved cell recovery rates using BEACON. a)** Joint distribution of barcode sample labels among the set of cells with concordance between cell line predictions using BEACON, Demultiplex, and transcriptome clusters. A concordance rate of approximately 99% is observed between Demultiplex and BEACON within individual sample labels. **b)** EMT gene scores as a function of pseudotime as reproduced using scripts from the original publication. **c-d)** EMT gene scores using the subset of cells with concordant transcriptome-based cell line assignments that are either shared between Demultiplex and BEACON, or identified by BEACON alone. **e)** UMAP representation among the cells in Fig. 3d of the main text. Cells are colored according to the subset as defined in the main text. **f)** Distribution of cell subsets within each of the annotated transcriptome clusters in **e)**. **g)** GSEA of the Hallmark EMT gene set within the ranked list of differentially expressed genes in each of the indicated time series. Here, we included the 10,000 most abundant genes within each time series to perform differential gene expression. **h)** Upset plot of all cells with concordant cell line assignments using various demultiplexing methods. **i)** Number of cells having concordant assignments with transcriptome-based clusters shared with at least 1 other demultiplexing method, stratified by different methods. **j)** Number of cells with concordant transcriptome-based cluster assignments that are uniquely identified by each of the indicated methods. **k)** Distribution of -log(FDR) for all genes in the data set, calculated using cells identified by the specified methods. FDRs were calculated for all twelve of the time series. Presented results include genes induced or repressed by ligand treatment.

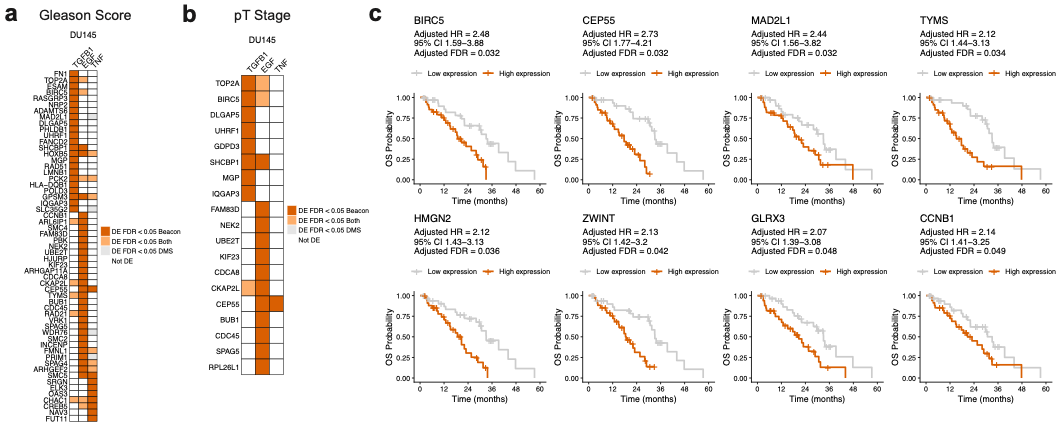

**Supplementary Figure 9. Genes nominated by BEACON that are associated with aggressive clinical features of prostate cancer. a)** Genes that are identified as differentially expressed in DU145 cells in any of the three ligand treatments using BEACON alone, as compared to Demultiplex-supervised (DMS), that additionally correlate with Gleason score in primary prostate adenocarcinoma tumors. **b)** Similar to **a)** but including genes that correlate with pathologic T-stage in primary prostate adenocarcinoma tumors. **c)** Genes identified as differentially expressed in DU145 cells in any of the three ligand treatments using BEACON alone that additionally associate with poor overall survival in a cohort with metastatic castrate resistant prostate cancer. Shown are Kaplan-Meier survival curves stratified by gene expression above and below the median.

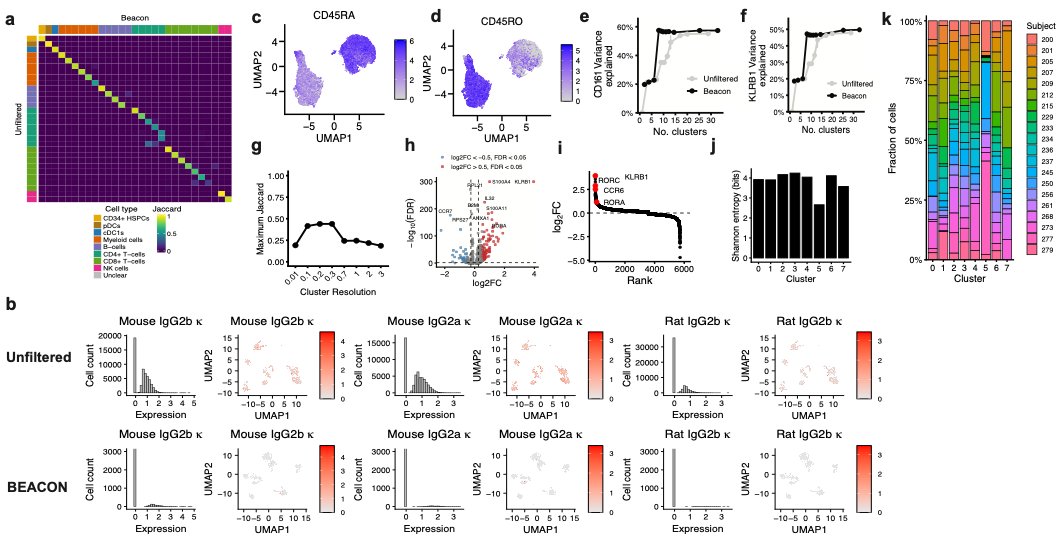

**Supplementary Figure 10. Application of BEACON to CITE-seq of human peripheral blood mononuclear cells. a)** Jaccard similarity between cell clusters identified using BEACON-filtered and unfiltered ADT data. Louvain clustering resolution parameter of 1 was used in Seurat for both sets of data. **b)** Mouse and rat isotype antibody controls without (top row) and with (bottom row) BEACON filtering. **c,d)** CD45RA and CD45RO ADT expression of CD4 T-cells as visualized on a UMAP generated from ADT marker expression. **e,f)** CD161 ADT and KLRB1 mRNA expression variance explained by clusters as a function of cluster number across a sweep of clustering parameter resolutions using BEACON-filtered and unfiltered ADT values. **g)** Maximum Jaccard similarity to the CD161-high cluster (cluster 3 in Fig. 4d of the main text) across a sweep of Louvain clustering resolution parameters, with clusters defined using the single-cell transcriptome data. **h)** Volcano plot of differentially expressed genes comparing cluster 3 to all CD4 T-cells using the Wilcoxon rank-sum test in Seurat. **i)** Rank abundance plot of log fold-change in expression of genes in cluster 3 against all other CD4 T-cells. **j,k)** Shannon entropy and relative donor representation within clusters defined in Fig. 4d of the main text.

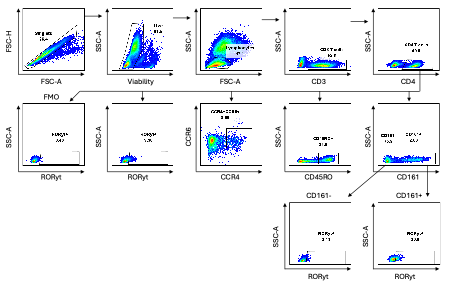

**Supplementary Figure 11. Flow cytometry of human peripheral blood mononuclear cells.** Representative gating strategy from a single donor for identification of CD3+, CD4+, CD45RO+, CD161+ and CD161−, CCR4+CCR6+, and RORγt+ and RORγt− T-lymphocyte populations.

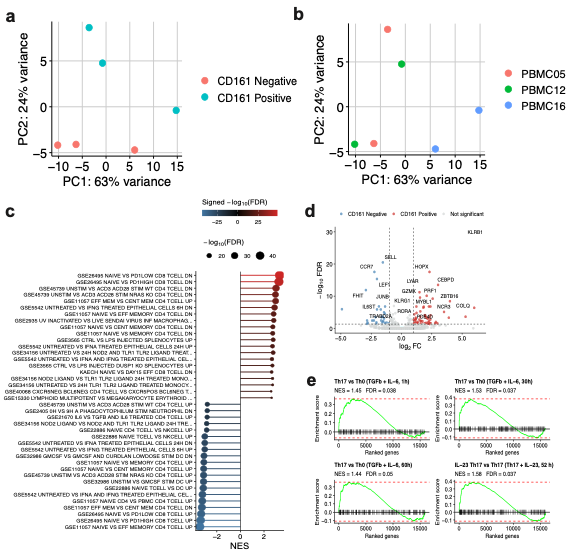

**Supplementary Figure 12. Bulk RNA sequencing of CD161-positive and CD161-negative cells within CD4-positive, CD45RO-positive T-cells of three healthy donors. a-b)** PCA of normalized expression values across the three donors. **c)** GSEA of genes ranked by DESeq2 Wald statistic using MSigDB immunologic signature gene sets, showing the 20 most positive and negative normalized enrichment scores. **d)** Volcano plot of differentially expressed genes in the CD161-positive vs. CD161-negative populations. **e)** GSEA enrichment of gene sets from the study of Yosef et al. derived from treatment conditions promoting Th17 CD4 T-cell differentiation.

| Markers | Fluorophore | Clone | Catalog/company | Dilution |
| --- | --- | --- | --- | --- |
| CD3 | BUV395 | UCHT1 | 563546/BD Biosciences | 1 to 1000 |
| CD4 | cFluor b548 | SK3 | 612889/Cytek Biosciences | 1 to 500 |
| CD161 | PE | HP-3G10 | 339903/BioLegend | 1 to 20 |
| CD45RO | PE-Cy7 | UCHL1 | 304229/BioLegend | 1 to 20 |
| CCR4 | APC | L291H4 | 359407/BioLegend | 1 to 20 |
| CCR6 | Alexa Fluor 700 | G034E3 | 353433/BioLegend | 1 to 20 |
| LIVE/DEAD Aqua | Aqua viability dye | N/A | L34966/Thermofisher | 1 to 1000 |
| RORγt | PE-CF594 | Q21-559 | 567532/BD Biosciences | 1 to 20 |

**Supplementary Table 1. List of flow cytometry markers used in the study.**
